# Construction of a reference genome for *Starmerella batistae* and annotation of *Starmerella* species reveal a close evolutionary relationship with *Schizosaccharomyces pombe* and suggest an alternative pathway for sophorolipid production

**DOI:** 10.1101/2025.09.24.678367

**Authors:** Soukaina Timouma, Alistair Hanak, Laura Natalia Balarezo Cisneros, Ian Donaldson, Fernando Valle, Daniela Delneri

**Affiliations:** Manchester Institute of Biotechnology, University of Manchester, Manchester, UK; University of Manchester Genomics Core Facility; BP Biosciences Center, San Diego, CA, USA

## Abstract

The *Starmerella* clade is known for displaying osmotolerant and acidophilic traits from their association with bees. Several species in this genus can produce sophorolipids, which are commercially produced as biosurfactants. Here, we isolated a yeast contaminant from the laboratory environment, identified as *Starmerella batistae,* able to thrive under low pH, including high concentrations of lactic acid, and relative high temperatures. We have sequenced and conducted a *de novo* genome assembly in three chromosomes and a mitochondrial genome for *S. batistae* (ca 9.3 Mb). Based on this reference genome we functionally annotated 29 different *Starmerella* species, using the publicly available sequences. Phylogenetic analysis across different yeast clades revealed a close relationship of *Starmerella* species with *Schizosaccharomyces* yeasts. Fifteen genes were uniquely shared between *Sz. pombe* and *S. batistae*, of which twelve were involved in cell morphology. Interestingly, the shape of *S. batistae* cells is elongated rather than round, similar to the *Sz. pombe*. Additionally, we found that all the *Starmerella* strains capable of producing sophorolipids shared a last common ancestor. Such clustering can help identify other sophorolipid-producing *Starmerella* yeasts that have not yet been characterised. We did not find the one-to-one orthologs of *S. bombicola* sophorolipid pathway in any of the *Starmerella* sp. with the exception of *S. kuoi*, *S. powellii* and *S. floricola*. In *S. etchelsii,* the antisense and telomeric pair UGTA1/CYP52M1 was found to be structurally conserved although not functionally. These findings support the notion that alternative pathways for the production of sophorolipids have evolved in different *Starmerella* lineages.

## Introduction

*Starmerella* yeasts are associated with insects and flowering plants and are typically well adapted for growth in high sugar environments (Spencer *et al.,* 1970; Rosa *et al.,* 2024). The genus *Starmerella* was first introduced to accommodate the haploid sexual stage of *Starmerella bombicola* (Rosa and Lachance, 1998). However, more recently, the widespread availability of DNA sequences has allowed for the robust establishment of the genus and the reassignment of 25 species previously assigned to the genus *Candida* (Santos *et al.,* 2018). *Starmerella batistae*, previously *Candida batistae*, was isolated in Belo Horizonte (Brazil) from the larval provisions of solitary digger bees *Diadasina distincta* and *Ptilotrix plumate*, larvae and pupae (Rosa *et al.,* 1999). Bee nests usually host a diverse community of microorganisms, including bacteria, moulds and yeasts. The yeasts are thought to have a key role in the conversion of pollen to beebread. Specifically, *S. batistae* and *Mucor* sp. yeasts in the community were found to have a role in the maturation of beebread (Rosa *et al.,* 1999). Several species within the genus maintain a stable haploid state (Rosa and Lachance, 1998), present with an elongated shape and reproduce asexually by budding either bilaterally or multilaterally (Melo *et. al,* 2014). In *S. bombicola*, it has been reported that single ascospores are formed, and cells are capable of conjugating with cells of the opposite mating type (Rosa and Lachance, 1998).

*Starmerella sp.* have gained considerable popularity and attention in various industrial sectors due to their unique metabolic capabilities and biotechnological applications. It is established that *Starmerella* yeasts harbour unique traits such as fructophily and the production of sophorolipids, which are one of the most commercialised glycolipid bio-surfactants (Kobayashi et al., 2024; Gonçalves *et al.,* 2020). Contrary to the petrochemical-based surfactants, bio-surfactants such as sophorolipids possess non or low ecotoxicity (Franzetti *et al.,* 2012), high biodegradability (Lima *et al.,* 2011), good biocompatibility (Lourith *et al.,* 2009) and excellent surfactant activity (Henkel *et al.,* 2012). *S. bombicola* is currently used industrially and shows the highest productivity of sophorolipids (Kim *et al.,* 2021).

In industrial fermentations, tolerance to low pH and high temperatures are widely desirable traits for production species. Significant energy costs are accrued through cooling of fermentations to counteract the heat released from metabolic reactions (Curran et al., 1999). Sophorolipid fermentations are also typically carried out at 25°C, aligning with the optimum growth temperature of *S. bombicola* (Van Bogaert *et al.,* 2007). However, running fermentations at higher temperatures improves heat transfer efficiency and allow modest cooling set-ups, which can translate to lower operating costs. Tolerance to acid conditions is also important in industrial fermentations since organic acids accumulate as by-products or in the case of lactic acid and citric acid, as the desired product (Martinez et al., 2013). Amelioration of media by adding base to reduce the acidity so that production species can maintain growth can make up a significant cost of production and cause downstream processing issues (Chen and Neilsen, 2016). We found that *Starmerella batistae* is capable of growing in such stress conditions, making it an interesting candidate for industrial fermentations.

Sophorolipids are composed by a disaccharide (β-1,2-linked glucose residues) and linked with a hydroxylated fatty acid via one terminal (ω position) or subterminal (ω-1 position). *S. bombicola* has been shown to produce sophorolipids primarily of the ω-1– hydroxylated fatty acid form while *S. batistae* primarily produces ω–hydroxylated fatty acid form. (Konishi *et al*., 2008). Sophorolipids with a fatty acid hydroxylated at the ω- position are known to be difficult to prepare by chemical synthesis, and are of interest for the polymer, fragrance and perfume industries (Kim *et al.,* 2021). Both *S. bombicola* and *S. batistae* are reported to produce a mixture of lactonic and acidic forms of sophorolipids, with the lactonic form (Kobayashi *et al.,* 2024) and acidic form (Konishi et al., 2008; Kim *et al.,* 2021) being the predominant component, respectively. Lactonic and acidic forms of sophorolipids exhibit distinct biological and physico-chemical characteristics (Van Bogaert *et al.,* 2007). The hydrophilic/lipophilic balance, foam generation capacity, and antimicrobial properties of these bio-surfactants are significantly influenced by the extent of lactone formation. Generally, lactonic sophorolipids demonstrate greater antimicrobial efficacy, while the acidic variants exhibit enhanced surface tension reduction foam production and solubility (Shah *et al.,* 2005). Although *S. bombicola* is used industrially and shows the highest productivity of sophorolipids (Kim *et al.,* 2021), the sophorolipids derived from *S. batistae* present potential for novel applications distinct from those associated with sophorolipids generated by *S. bombicola*.

Here, we have isolated a lab contaminant displaying low pH and high temperature tolerance and identified it as a genetic variant of the type strain *S. batistae* CBS 8550. A *de novo* telomere-to-telomere genome assembly was produced, including the mitochondrion, and structural and functional annotation of *S. batistae* alongside 29 assembled, publicly available, *Starmerella* yeast species, were carried out. We identified genus-specific and species-specific genes, and genes uniquely shared with *Sz. pombe* involved in cell morphology. All the known sophorolipid producing species clustered together and shared a close last common ancestor. Intriguingly, orthologs of *S. bombicola* genes of the sophorolipid biosynthetic pathway were not found in all of the other species, suggesting the existence of alternative pathways for their production, possibly linked to the type of sophorolipids produced.

## Results and Discussion

### Phenotypic characteristics of *S. batistae*

A strain of *S. batistae* was isolated as a contaminant in our lab and named *Starmerella batistae* var. SB001. Sanger sequencing of the internal transcribed spacer (ITS1) and D1/D2 domains of ribosomal DNA (rDNA), followed by BLAST alignment against the NCBI database revealed 100% identity of both genes with *Starmerella batistae* type strain, CBS 8550, and is closely related to *Starmerella riodocensis, Starmerella kuoi* and *Starmerella bombicola*, as shown in the phylogenetic tree using the ITS1 sequence (Figure 1). Firstly, we have carried out a standard phenotypic characterisation and compared it with the type strain *S. batistae* CBS8550 isolated by Rosa *et al* 1998. Colony morphology and assimilation of several sugars were consistent with previous descriptions and both strains of *S. batistae* were able to grow on glycerol as sole carbon source, hence suggesting the presence of functional mitochondrion (Supplementary Figure 1). This was also supported by the DAPI staining to visualise the nuclear and mitochondrial DNA content, confirming that *S. batistae* SB001 *is* rho+ (Supplementary Figure 2).

**Figure 1.**
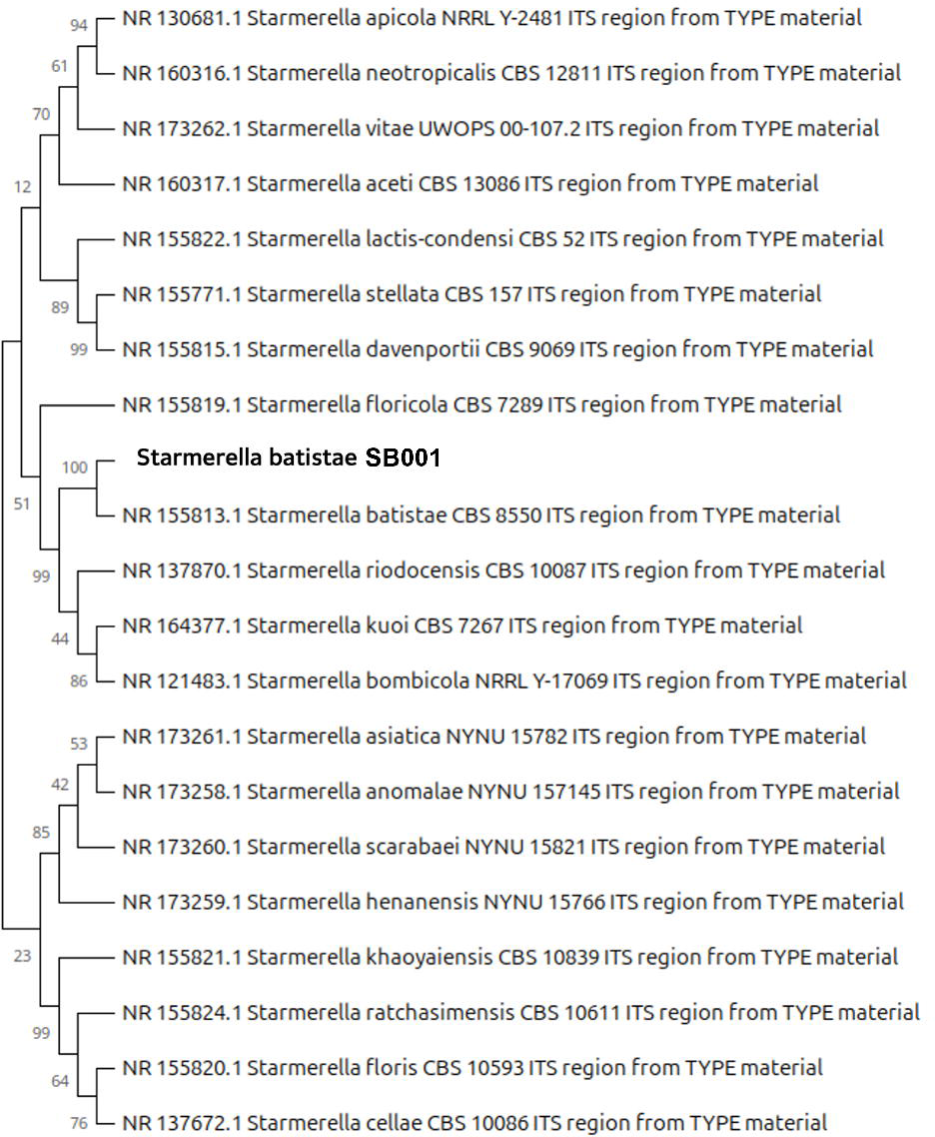

Variations in the growth profile of SB001 and CBS8550 were observed at different temperatures, with SB001 showing a higher growth rate at 25°C compared to CBS8550 (Supplementary Figure 3, panel A), while at 32°C and 36°C CBS8550 grew better than SB001 (Supplementary Figure 3, panels A and B). At 43°C both strains struggled to grow (Supplementary Figure 3 panel B). *S. batistae* SB001 is also able to thrive at low pH (*i.e.* pH 2.0) in a different range of temperatures (Supplementary Figure 4 panels A and B) and can efficiently grow in the presence of 60g.l^-1^ of lactic acid at 37°C (Supplementary Figure 4, panel C).

### Genome sequencing and telomere-to-telomere *de novo* assembly of *S. batistae*

Genome sequencing of *S. batistae* SB001 was performed using HiFi read data derived from single-molecule real-time (SMRT) technology from Pacific Biosciences (PacBio). The IPA program assembled the HiFi reads data set into 3 gap-less chromosomes bounded by telomeric repeats, totalling *ca.* 9.3 Mb in length, as well as one mitochondrial genome of 38 kb. The chromosome sizes were 5.22 Mbp, 2.18 Mbp and 1.86 Mbp (Supplementary Table 1). Given no haplotig was detected, the *S. batistae* SB001 isolate is likely to be haploid. The sequencing reads were mapped to the genome assembly using Minimap2 with parameters tailored for PacBio sequencing reads and the variant calling analysis was done using DeepVariant (Poplin *et al.,* 2018). As result, only 12 biallelic SNPs were found across the whole genome, supporting the fact that this isolate is haploid (*i.e.* such a low number of SNPs are likely to be sequencing errors; Supplementary Figure 5). The related species *S. bombicola* is known to exist in both diploid and haploid forms (Rosa and Lachance, 1998).

To validate the mating type of *S. batistae* SB001, we looked at the expression of the **a**/α pheromone receptors. The expression of the a-factor pheromone receptor *STE3* (SBAT_0B07760), which is only present in MATα cells (Sprague et al., 1983), is negligible in *S. batistae*, while the α-factor pheromone receptor *STE2* (SBAT_0C05970), which is only present in MAT**a** type cells (De Segni *et al.,* 2011), was clearly expressed (Supplementary Figure 6), hence confirming *S. batistae* SB001 as haploid MAT**a**.

Overall, given the biotechnological importance of *Starmerella* sp., the high-quality assembly of the *S. batistae* genome will be crucial for targeted genetic manipulation of this yeast. Additionally, the haploid state of *S. batistae* will allow for more straightforward engineering and for potential hybridisation.

### *S. batistae* structural and functional annotation reveals a low level of redundancy

The structural annotation of *S. batistae* genome was carried out using YGAP (Proux-Wéra *et al.,* 2012) which identified a total of 4480 structural elements (2248 on strand + and 2232 on strand -), including 4379 protein coding genes, 101 tRNAs and interestingly only 3 TY-like retrotransposons (Figure 2). The output of YGAP for the *S. batistae* predicted genes used the following nomenclature: SBAT for *S. batistae,* and consecutive alphabet letters were used for the chromosome number. Functional annotation was conducted by identifying one-to-one orthologs between *S. batistae* proteins and well-annotated model organism proteomes including *S. cerevisiae*, *C. glabrata*, *C. albicans*, *Y. lipolytica* and *S. pombe.* Out of the 4379 *S. batistae* proteins, 3906 (*ca.* 90%) were successfully functionally annotated as they had a one-to-one ortholog in at least one of the 5 yeast model organisms (Supplementary Table 2). The mitochondrial DNA possesses 10 protein coding genes, including one-to-one orthologs of *S. cerevisiae VAR1, OLI1, ATP6, COB, COX2, NAD4* and *AI2*, as well as 16 tRNAs.

**Figure 2.**
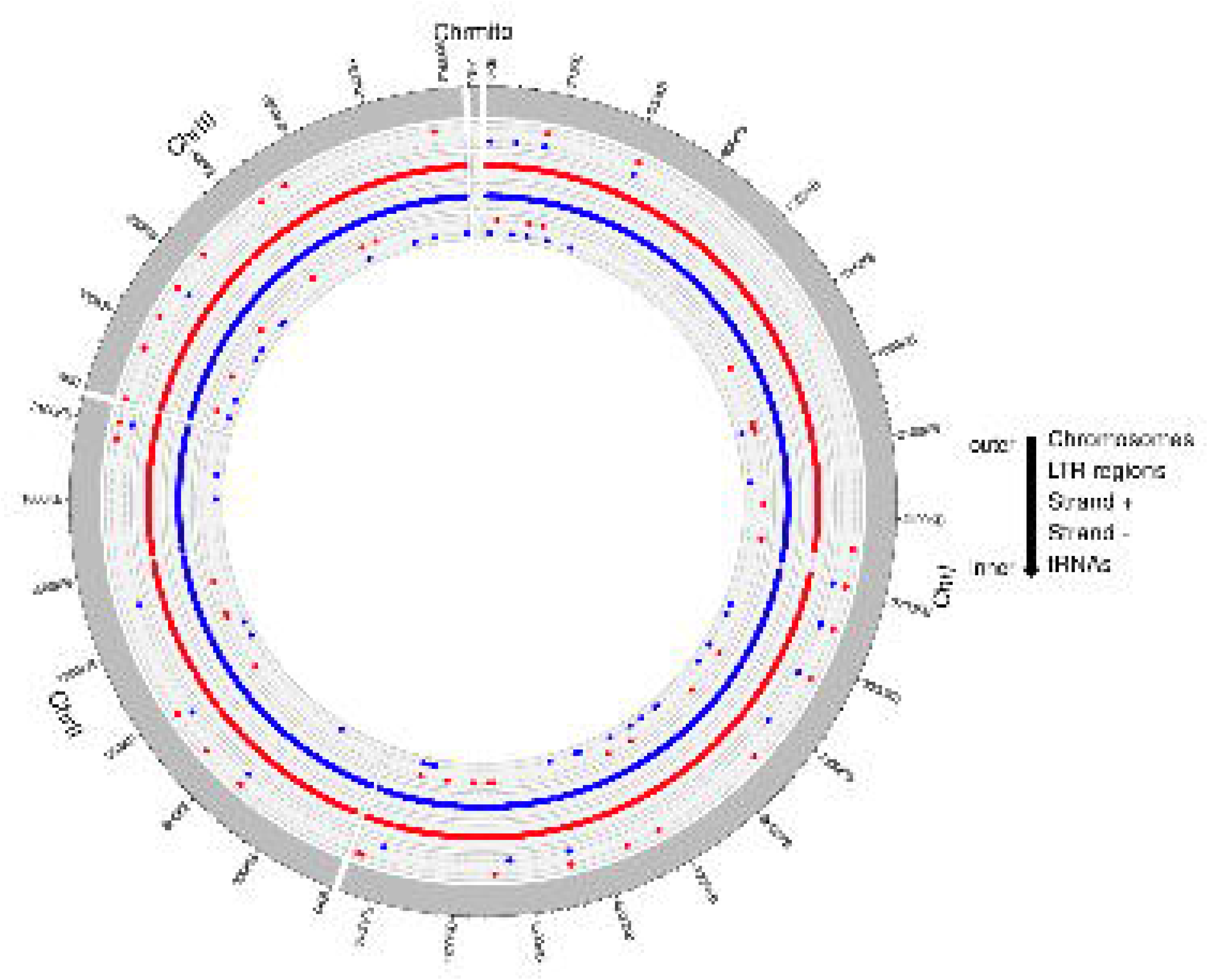

The genetic redundancy within *S. batistae* genome was investigated by predicting group of homologues including paralogs using HybridMine (Timouma *et al.,* 2020). As result, 21 group of homologous genes were identified (93 paralogs in total), of which 4 groups of homologs contain a young paralog sharing 100% identity. Moreover, 72 LTR retrotransposons were detected using LTR_retriever (Figure 2; Supplementary Table 3). TY-like elements have the ability to trigger non-allelic homologous recombination between LTRs, which can lead to segmental, tandem or partial duplication of genes (Coughlan and Wolfe, 2018). We found that 11 out of 21 homologues group, had paralogs genes located in LTR retrotransposon regions and hence may have arisen from segmental duplication via LTR recombination (Supplementary Table 3). Overall, the level of redundancy in *S. batistae* SB001 is low and may reflects the low number of TY-like transposons and solo LTRs. In fact, in comparison, 124 tandem duplicates have been identified in *C. albicans* (Fitzpatrick *et al.,* 2010), while for the post-WGD yeast *S. cerevisiae* 1356 duplicate genes have been detected (Ames et al., 2010).

### Identification of variation between *S. batistae* SB001 and CBS 8550

*S. batistae* SB001 and the type strain CBS8550 were both sequenced by illuminaNovaseq6000 and compared against the newly assembled reference genome of SB001. Short variants (indels and SNPs) were identified between the two genomes revealing 23 unique variations, 4 of which were located in coding regions (Supplementary Table 4). These short variants were verified and confirmed by sanger sequencing and included: *i. SBAT0C00870* (*SYT1* orthologs in *S. cerevisiae*), a guanine nucleotide exchange factor involved in exocytosis; *ii. SBAT0C06340* with a predicted function in RNA splicing; *iii. SBAT0A02050* (*ENV9* orthologs in *S. cerevisiae*), an oxidoreductase involved in vacuolar morphology; and *iv. SBAT0A15890* with a predicted function in RNA metabolism (Supplementary Table 4). These genetic variants may alter the functionality of the genes and be responsible for the difference in growth phenotype observed at different temperatures (Supplementary Figure 3).

Overall, the two strains showed a high degree of similarity and can be confirmed as variants of the *S. batistae* type strain.

### Genome annotation of *Starmerella sp*ecies and identification of the core genome

We have accessed the genome assemblies 29 *Starmerella* yeasts available on NCBI and carried out the structural and functional annotation of their genome (Supplementary Table 5). Such data analysis enabled us to determine the coregenome (*i.e.* the set of genes present in all species analysed). The accessory-genome (*i.e.* specific to one or a group of species) was difficult to annotate due to lack of functional profiling of the different *Starmerella* species. The core genome included 1791 protein coding genes (Supplementary Figure 7). Given the fact that the majority of the *Starmerella* yeasts have an incomplete or poorly assembled sequences, with contigs ranging from 4 to 5488 (Supplementary Table 5), it is very likely that this number of core genes is under-estimated. Out of the 1791 genes in the *Starmerella* coregenome, 1687 share a one-to-one ortholog with *S. cerevisiae* (Supplementary Table 2). As expected, the gene ontology enrichment analysis reveals significant over-representation (p-adjusted value < 0.001) of fundamental biological processes such as cellular and metabolic activities, including regulation of chromosome organisation, cell cycle, or specific pathways like ribosome biogenesis and RNA processing, highlighting the essential functions supported by the core genome (Supplementary Figure 8).

### Phylogenetic analysis of *Starmerella sp*. yeasts reveals a close relationship with *Schizosaccharomyces* yeasts

One-to-one orthologs between *S. batistae* and five model organisms (*i.e. Saccharomyces cerevisiae*, *Candida glabrata*, *Candida albicans*, *Yarrowia lipolytica* and *Schizosaccharomyces pombe*), were used for functional annotation. This analysis highlighted 10% of genes lacking annotation that could be involved in specific functions, such as growth at high temperature and low pH adaptations (Supplementary Figure 4). To identify the latter, we also predicted one-to-one orthologs between *S. batistae* and five acidophilic yeasts (*Kluyveromyces lactis, Maudiozyma exigua, Maudiozyma barnetti, Maudiozyma bulderi and Kluyveromyces marxianus*), as well as three phylogenetically close relatives (*Wickerhamiella sorbophila, Saprochaete ingens,* and *Sugiyamella lignohabitans*). This analysis revealed 83 *S. batistae* proteins, lacking a one-to-one ortholog in the five model organisms, that had orthologs in at least one of the low pH-tolerant yeast species (Supplementary Table 2). In total, 2,019 *S. batistae* proteins were conserved across all 13 yeast species included in the orthology analysis (*i.e.*, 5 model organisms, 5 acidophilic species, and 3 closely related yeasts; see Figure 3). As expected, *W. sorbophila* shared the highest number of one-to-one orthologs with *S. batistae* (Supplementary Table 2). Furthermore, we identified 242 proteins unique to *S. batistae*, which may include strain-specific or lineage-specific proteins (Figure 3). Interestingly, 15 proteins were found to be uniquely shared between *S. batistae* and *S. pombe* (Figure 3). Enrichment analysis indicated that these proteins are predominantly associated with cell morphology (adjusted p < 0.001; Figure 4; Supplementary Table 6).

**Figure 3.**
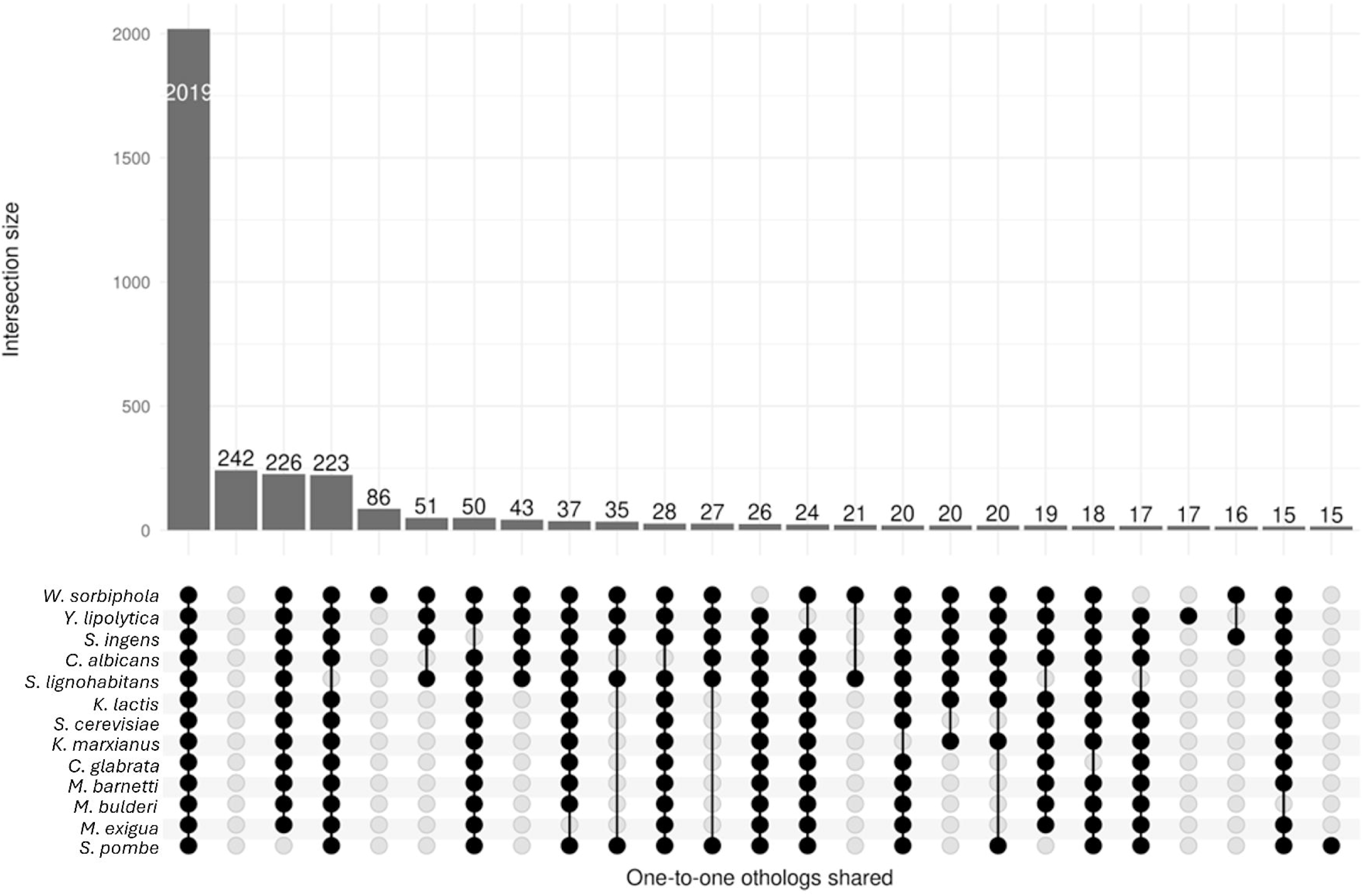

**Figure 4.**
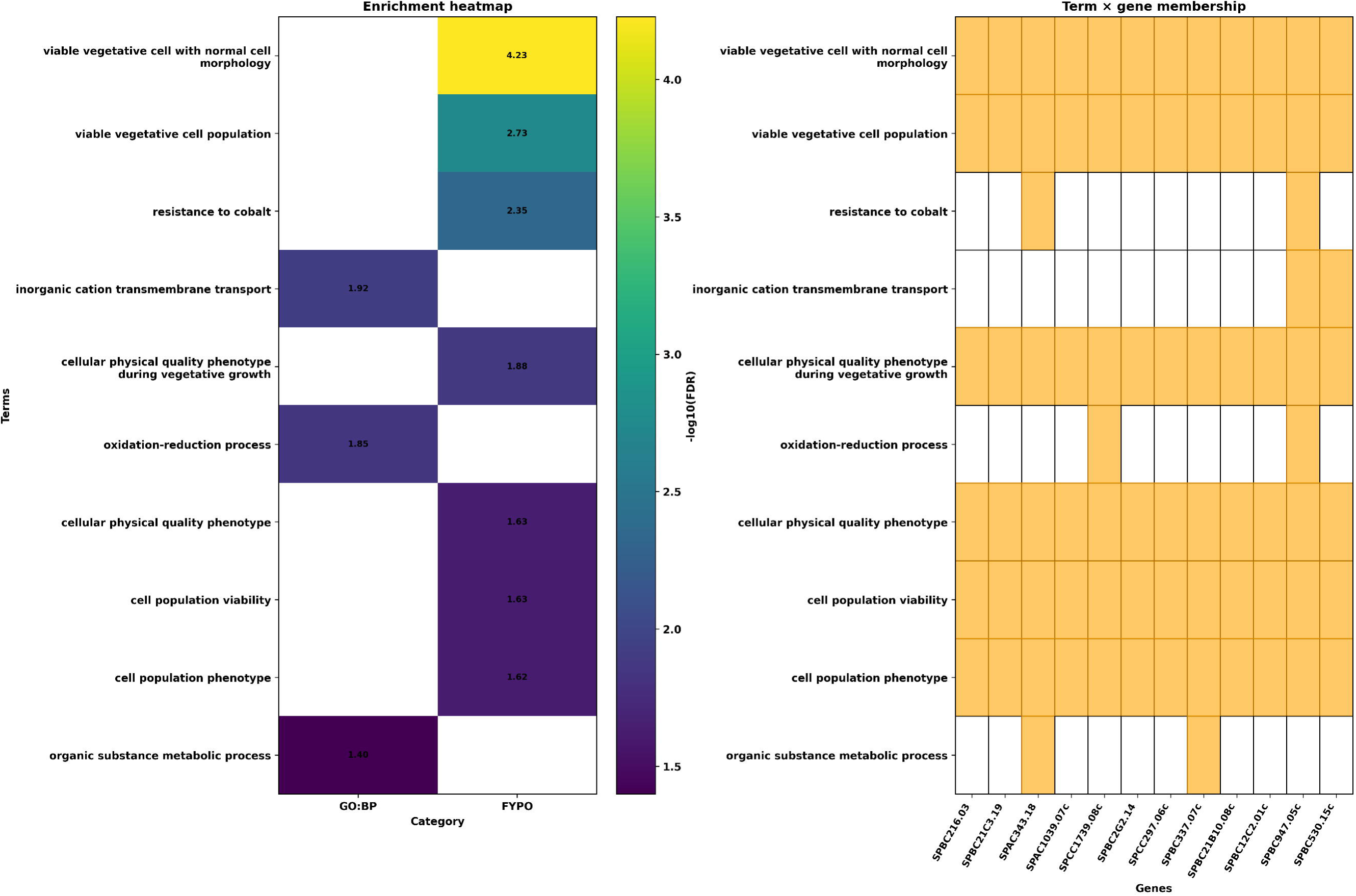

There are 17 proteins that were found to be unique with *Y. lipolytica* (Figure 3; Supplementary Table 6). No significant process enrichment was found; however, they possess diverse functional roles in cellular integrity, gene regulation, mRNA metabolism, lipid metabolism, and environmental adaptability (Supplementary Table 6).

A first phylogenetic tree was constructed based on the 2,019 shared orthologous proteins (Figure 5, panel A). The resulting phylogeny demonstrated well-supported distinct clades: *Starmerella, Schizosaccharomyces, Candida, Kluyveromyces,* and *Maudiozyma/Kazachstania*. Notably, *S. batistae* grouped most closely with the *Schizosaccharomyces* lineage, suggesting an unexpected evolutionary relationship. Additionally, *Sugiyamaella lignohabitans, Schizosaccharomyces,* and *Starmerella* species were the only taxa examined that possessed three chromosomes plus mitochondrial DNA, whereas *Saprochaete ingens* contained five chromosomes (Hodorová et al., 2019). The pre-whole genome duplication clades *Sugiyamaella*, *Saprochaete* and *Starmerella* all share the last common ancestor with to the *Schizosaccharomyces* clade.

**Figure 5.**
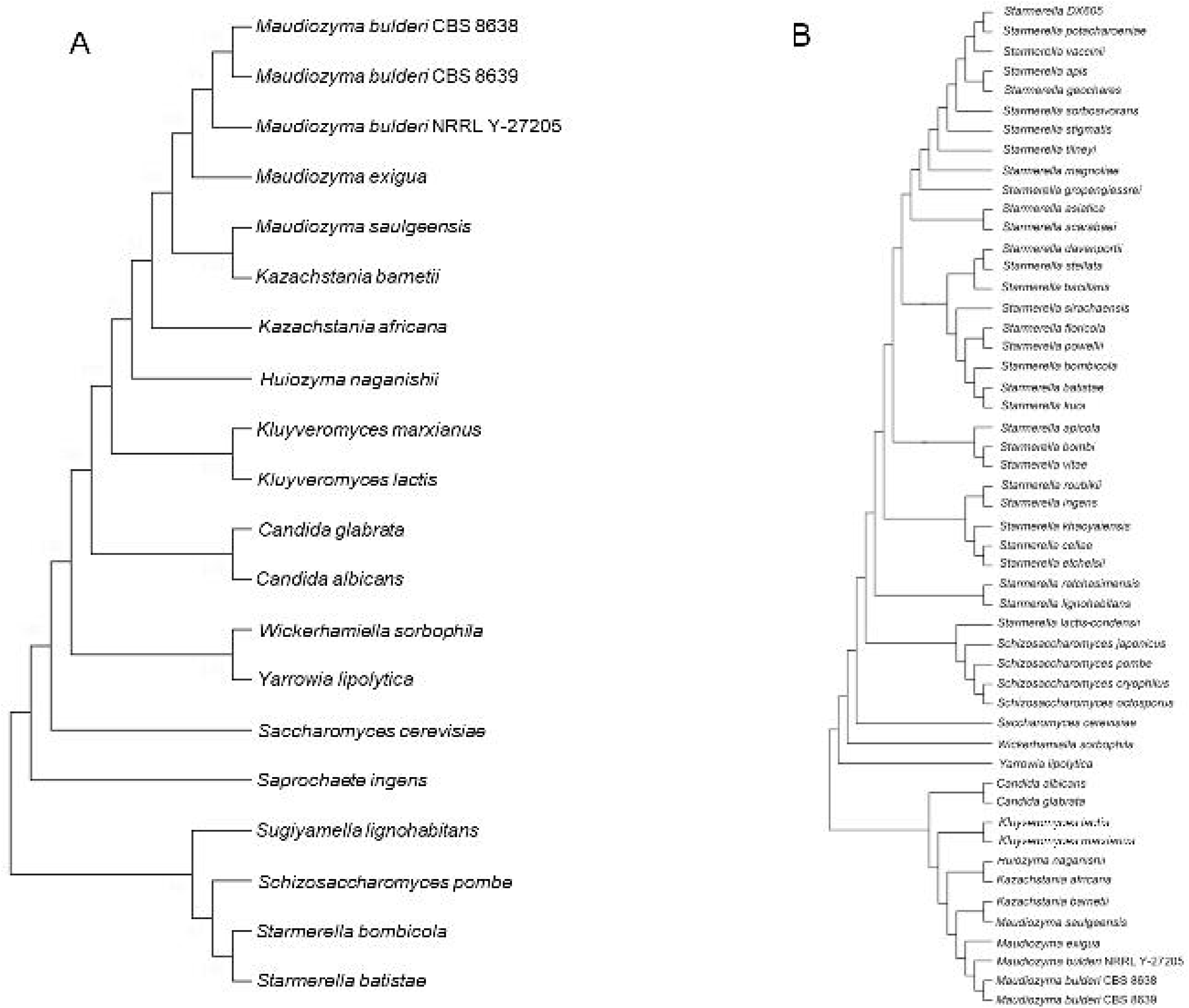

A second tree was constructed using 1,288 core proteins shared with 3 additional yeasts from the *Schizosaccharomyces* lineage and all the *Starmerella* yeasts with sequenced genome (Figure 5, panel B). Both trees yielded a similar topology, with the *Starmerella* clade clustering close to *Schizosaccharomyces* clade, supporting the robustness of the phylogenetic inference.

*Starmerella lactis condensi* was found to be phylogenetically closer to the *Schizosaccharomyces* clade than the *Starmerella* clade. Given the ITS tree place this species closest to *Starmerella stellata* (Figure 1), it is possible that the quality of genome assembly is the reason for this taxonomical anomaly.

The phylogenetic proximity with *Schizosaccharomyces* clade, alongside with the uniquely shared genetic content with *Sz. pombe* (Figure 3) raises intriguing questions, including the possibility of ancient introgression events, which warrant further investigation in the context of *S. batistae*’s evolutionary history. It worth noting that both *Schizosaccharomyces* and many *Starmerella* yeasts naturally present with an elongated shape (Rosa and Lachance, 1998), and, interestingly, *S. acetii*, for which there is no genome available, reproduce asexually by bilateral budding (Melo *et. al,* 2014).

### Comparative genomics of sophorolipid production genes suggest alternative pathways in *Starmerella species*

In total, 32 *Starmerella* species have been sequenced (Shen *et al*., 2018; Opulente *et al*., 2024), of which several are known to produce sophorolipids (Konishi *et al.,* 2008; Kurtzman et al, 2010; Kurtzman 2012). Production and characterisation of the sophorolipids have been extensively studied in *S. bombicola.* These molecules were also detected in *S. riodocensis, S. apicola, S. batistae, S. kuoi,* and *S. stellata*, but with limited biochemical studies up to date. It is known that accurate characterisation of the diverse sophorolipid structures produced by different species is not straightforward and the availability of standards for chemical analysis is poor (Ingham et al., 2024). As a result, sophorolipid mixtures produced by different species are likely not fully characterised.

*S. bombicola* is the primary species utilised in industry due to its apparently highest production yield. It is known that the *Starmerella* species produce various classes of sophorolipids, primarily differentiated into acidic and lactonic forms, as well as ω and ω-1 types with different levels of acetylation (Konishi et al., 2008). Some species are reported to produce a mixture of sophorolipids in different proportion and more unusual forms including sophorolipid polymers such as dimeric and trimeric sophorolipids (Kurtzman., 2012).

The biosynthetic pathway for sophorolipids production in *S. bombicola* is the best described and starts with the import of fatty acids to the cell, which proceed through hydroxylation by a cytochrome P450 52-M1 (encoded by the *cyp52M1* gene), then two glycosyltransferase reactions (encoded by *UGTA1* and *UGTB1* genes), ultimately resulting in the formation of sophorolipids. These molecules can be acetylated by an acetyl transferase (encoded by the *AT* gene), and secreted by a multi-drug transporter (encoded by the *MDR* gene) before being lactonized by a lactone esterase (encoded by the *SBLE* gene; Figure 6, panel A). As control for the sophorolipid pathway research, we identified DNA and protein matches of the published sequences against the genome assembly of *S. bombicola*. As expected, all genes were found with 100% identity apart from *AT* (Figure 6, panel B). This was because the available sequence and assembly used (GCA_001599315.1) is from a mutant for *AT* (*S. bombicola* Δ*AT*; Saerens et al., 2011). Next, this *S. bombicola* sophorolipid production pathway was searched in all the *Starmerella* yeasts annotated using protein sequences.

**Figure 6.**
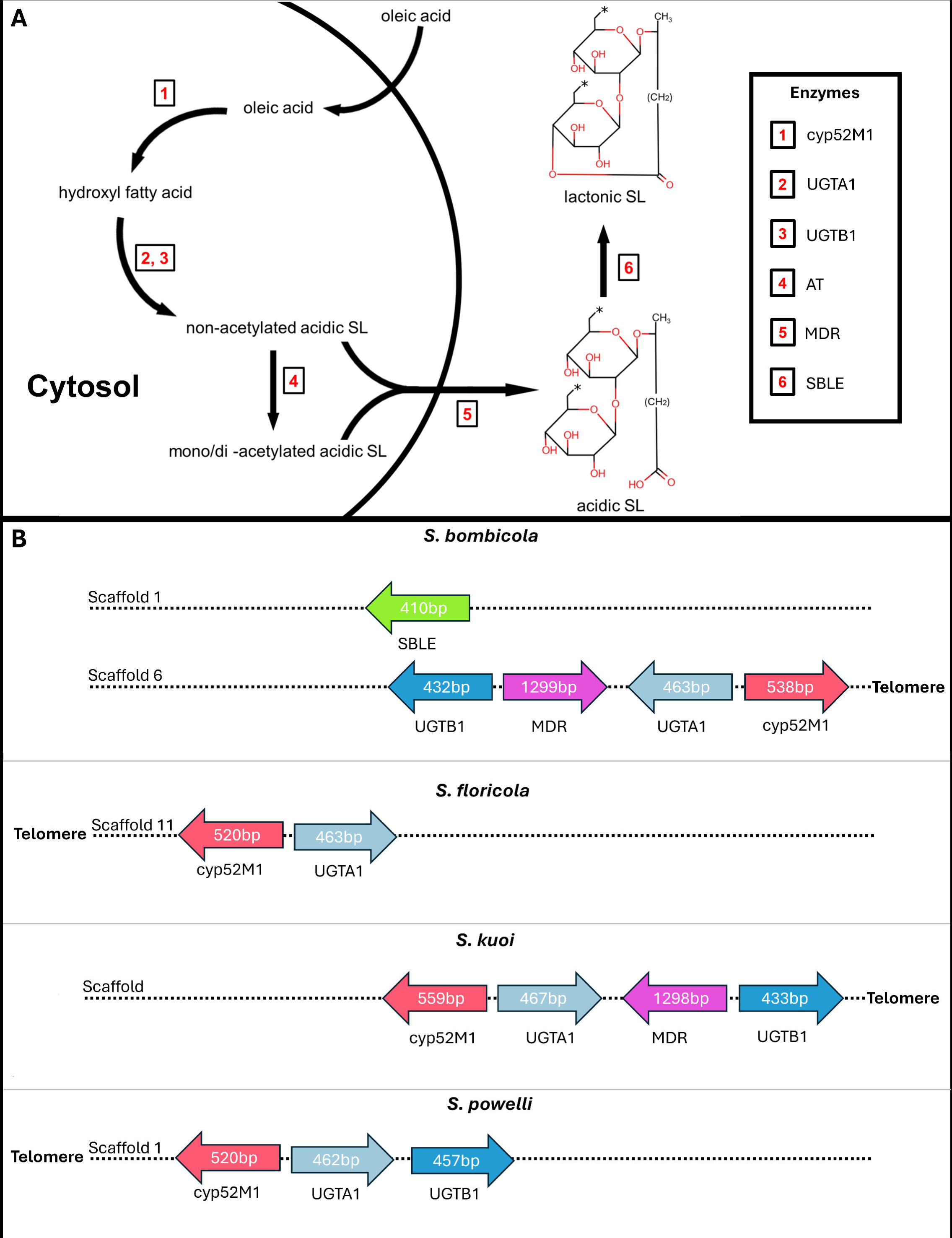

Species that are known to produce sophorolipids clustered together based on shared genome and identifies species that have not yet been characterised such as the highly acidophilic yeast *S. davenportii* (Supplementary Figure 9). These species, namely *S. vitae*, *S. bombi*, *S. apicola*, *S. kuoi*, *S. batistae*, *S. powellii*, *S. floricola*, *S. sirachaensis*, *S. bacillaris*, *S. stellata* and *S. davenportii* might have their last common ancestor possessing the ability to produce sophorolipids. Out of this cluster, *S. kuoi*, *S. stellata*, *S. apicola* have been described as producing mixtures of acidic and lactonic sophorolipids.

Based on one-to-one orthology search between the *S. bombicola* proteome versus all the *Starmerella* yeasts pan-proteome of this study, we could not find any one-to-one orthologs of the *S. bombicola* sophorolipid pathway in any *Starmerella* yeats including *S. batistae*, except for 3 species (Supplementary Table 7), where a limited number of orthologs were found: *S. floricola* shared *cyp52M1* and *UGTA1* (Figure 6, panel B), *S. kuoi* shared *cyp52M1*, *UGTA1*, *UGTB1* and MDR (Figure 6 panel B) and *S. powelli* shared *cyp52M1*, *UGTA1* and *UGTB1*. Notably, *UGTA1* and *cyp52M1* consistently conserved their antisense arrangement and structural co-location similarly to *MDR* and *UGTB1*. In *S. kuoi* the pairs of genes *UGTA1/cypM1* and *MDR/UGTB1* were inverted compared to the other species (Figure 6 panel B, Supplementary Table 7). *S. floricola*, *S. powelli* and *S. kuoi* are also the closest phylogenetically to *S. bombicola*. Of these species *S. kuoi* was found to produce a significant amount of sophorolipids, primarily in the acidic form (Kurtzmann *et al*, 2010), which is consistent with our annotation data which suggests the lack of lactone esterase in its genome. *S. floricola* produces only a small quantity of sophorolipids, which remain uncharacterised (Kurtzmann *et al*, 2010), while there is no information in literature on their production in *S. powelli*.

For many of the *Starmerella* species, the lack of other one-to-one orthologs from the sophorolipid production pathway could be an artefact due to the poor-quality assemblies available. However, it is unlikely these orthologs were missed in our well assembled genome of *S. batistae,* consisting of 3 gap-less chromosomes plus a mitochondrial genome (Supplementary Table 5).

When we expanded our comparative analysis to homologs of the *S. bombicola* sophorolipid pathway that showed structural conservation (*i.e.* the antisense pair UGTA1/CYP52M1 colocalised with the antisense pair UGTB1/MDR1). As expected from the orthology analysis, structural matches occurred in*, S. floricola, S. kuoi, S. powellii,* additionally, we found one additional species, *S. etchellesii*, potentially possessing some genes of the *S. bombicola* described pathway (Supplementary Table 9). The colocalised antisense pair UGTA1/CYP52M1 in *S. etchellsii* (ETC_0A00120/SETC_0A00110, respectively) was found to be also telomeric (scaffold 1) as described in *S. bombicola*. Interestingly, these genes (ETC_0A00120/SETC_0A00110) were annotated by YGAP as sharing an ancestor with ATG26 (sterol glucosyltransferase)/DIM1 (dimethyladenosine transferase), respectively. This antisense pair is structurally conserved across nearly all *Starmerella* species, but it appears too diverged at the sequence level to yield direct matches with the *S. bombicola* UGTA1/CYP52M1 proteins. This suggests that the pair in *S. etchellsii* may represent either a paralogous remnant of the sophorolipid pathway, or a more ancient and conserved antisense arrangement that has functionally diverged in most *Starmerella* lineages. The other antisense structural pair UGTB1/MDR was not found in any of the *Starmerella* yeasts. In *S. etchellsii,* UGTB1 didn’t generate a hit, however an MDR homolog (SETC_0S00360, scaffold 19) sharing 76% positives (60% identity) was found. Here, it is possible that the lack of match is due to the assembly quality (Supplementary Table 5).

Interestingly, in *S. batistae*, although homologs with low sequence identity were detected for *UGTB1* (27.1% identity), *AT* (42.7% identity), *MDR* (62.7% identity), *SBLE* (44% identity) and *cyp52M1* (42.4% identity), the pathway was not structurally conserved (Supplementary Table 8, Supplementary Table 9). This data together with the overall low redundancy in *S. batistae* points towards a different enzymes for the production of these molecules. A recent study suggested a revised pathway for the sophorolipid production in *S. bombicola* (Roelants et al., 2024). In fact, *S. bombicola* Δ*SBLE* strains have been found to produce acetylated “bola” form sophorolipids instead of acidic sophorolipids and Δ*AT* strains were still capable of producting acetylated sophorolipds, although at a deceased level, suggesting other undescribed genes may be involved in the sophorolipid synthesis pathway (Roelants et al., 2024).

Overall, our work also supports the hypothesis that different *Starmerella* species use alternative synthesis genes and potentially alternative pathways for the production of sophorolipids. For example, *S. batistae,* despite the lack of shared orthologs for the *S. bombicola* sophorolipid production pathway, can still produce similar sophorolipids. Additionally, *S. kuoi* which shares the most sophorolipid synthesis orthologs with *S. bombicola* is reported to produce dimeric and trimeric sophorolipids not known to be produced by the latter, suggesting it possesses additional synthesis genes.

## Conclusions

*S. batistae,* possesses several desirable traits for use in industrial application, being able to thrive in high concentrations of inorganic and organic acids, high concentrations of sugar and is thermotolerant relative to other yeast species.

The ability of the sophorolipid-producing *S. batistae* to grow optimally at higher temperatures, as we show in this work, can increase the efficiency of cooling fermentations, which can significantly lower production costs. This is especially relevant in the production of sophorolipids where the cooling is required for *S. bombicola* optimal growth (Wongsirichot & Winterburn 2024).

Here, we have isolated a lab contaminant displaying low pH and high temperature tolerance and identified it as a genetic variant of the type strain *S. batistae* CBS 8550. We produced a reference genome for *Starmerella batistae*, including the mitochondrion, via *de novo* telomere-to-telomere genome assembly. We confirmed the mitochondrion is functional as this yeast was able to grow on glycerol as sole carbon source. Variant calling analysis shows this strain is haploid and the expression of the α-factor pheromone receptor *STE2* suggests that this *S. batistae* is Mat***a***. Structural annotation of *S. batistae* was carried out alongside with 29 *Starmerella* yeasts with publicly available sequences. Functional annotation was performed using well-annotated model organisms including *Saccharomyces cerevisiae*, *Candida glabrata*, *Candida albicans*, Y*arrowia lipolytica* and *Schizosaccharomyces pombe.* By comparing the proteome of all these yeasts, we identified genus specific and species specific genes. Phylogenetic analysis across different clades of yeast including *Starmerella, Wicherhamiella, Saprochaete, Sugiyamella, Maudiozyma, Kazachstania, Yarrowia, Candida* and *Schizosacchaomyces* revealed an unexpected close relationship between the *Starmerella* and *Schzosaccharomyces* clade. Strikingly, fifteen genes were found uniquely shared between *S. batistae* and *Sz. pombe*, and were phenotypically enriched in cell morphology processes (FYPO term), potentially underpinning the trait of elongated cell shape of these yeasts.

A phylogenetic tree, constructed using 1288 shared proteins, highlighted that the sophorolipid producing *Starmerella* yeasts share a close last common ancestor. This cluster could potentially reveal other *Starmerella* yeasts with the ability to produce sophorolipids that have not yet been characterised, such as *S. davenportii* which is known to grow at pH as low as 1.4 (Stratford et al., 2002). The sophorolipid biosynthetic pathway described in *S. bombicola* was absent in all *Starmerella* species examined, except for *S. kuoi, S. powellii,* and *S. floricola*, which also happen to be the closest phylogenetically to *S. bombicola*. In *S. etchelsii*, the antisense telomeric gene pair UGTA1/CYP52M1 showed structural conservation but were found too divergent at the sequence level. All together, these results suggest that distinct *Starmerella* lineages may have evolved alternative mechanisms for sophorolipid production, which can potentially reflect the different classes and modifications of these molecules.

## Methods

### Phenotypic analysis

Growth curves were obtained using Flurostar Omega plate readers (BMG Labtech, Allmendgrün, Germany). Cells were grown overnight in YPD at 25°C and then washed twice with distilled water. Wells containing 200µl of growth media were inoculated with a starting OD_600_ of 0.1. Growth media consisted of Synthetic Minimal Media (SD media) supplemented with 2% glucose and pH was modulated using H_2_SO_4._ Media containing lactic acid was buffered to pH 2.5 with 1M NaOH. Optical density was measured every 1000s with shaking for 80s before each measurement at a width of 3mm for 250 cycles (Approximately 69 hours). Plate reader assays were carried out at 25°C, 32°C, 36°C, 37°C and 43°C.

### DNA extraction and library preparation for long read next generation sequencing

For PacBio sequencing, the DNA was extracted from samples using the cetyl trimethyl ammonium bromide (CTAB) method (Wu et al., 2001). For the CTAB method, 1[mL of overnight culture was added to 50 mg of acid washed glass beads 425–600 µm (Sigma). 1 mL of CTAB extraction buffer was then added, and after incubation at 65 °C. RNAse A treatment was conducted by adding 2 µL of RNAse A 100 µg/mL (Qiagen, Germany) and incubating the samples at 37 °C for 15 min. After purification with phenol:chloroform:isoamyl alcohol (25:24:1) the DNA was eluted in 50[L of ultrapure distilled water (Invitrogen) and stored at 4 °C. The DNA quality was assessed using the NanoDrop LiTE Spectrophotometer (Thermofisher Scientific) to be within the quality specification range required by the PacBio protocol. The genomic DNA was adjusted to 10 ng/µL in 150[µL and sheared to ∼10 kilobase fragments using g-TUBES (Covaris) following the manufacturer’s instructions. The quality and size of DNA fragments were verified using Fragment Analyzer (Advanced Analytical Technologies) following the DNF-90 protocol. Samples were prepared for sequencing following the Express Template Prep Kit 2.0 protocol (Pacific Biosciences, USA), with multiplexing using the Barcoded Overhang Adapter kit 8A (Pacific Biosciences, USA). DNA libraries were sequenced using the SMRT Cell 1 M chips on the Pacific Biosciences Sequel system with 10 h data acquisition time.

### DNA extraction and library preparation for short read next generation sequencing

For next generation short read sequencing, genomic DNA from Strains CBS8550 and SB001 was extracted using a MasterPure DNA extraction kit (LGC Biosearch Technologies, UK) according to the manufacturer’s instructions. DNA quality and concentration was checked with a Qubit 4 Flurometer (ThermoFisher, USA) and sequenced on the Illumina No-vaSEQ6000 (Illumina Inc., USA) platform. Unmapped paired-reads of 159bp from Illumina NovaSeq6000 were checked using a quality control pipeline including FastQC v0.12.1 (Babraham Institute 2010) and FastQ Screen v0.15.3 (Wingett and Andrews 2018). Indexes for reference genomes were created for bwa-mem (Li and Durbin 2010) using ‘bwa index’ v 0.7.17. Reads were trimmed to remove adapter sequences or poor-quality reads using Trimmomatic v0.39 (Bolger, Lohse, and Usadel 2014); reads were truncated at a sliding 4bp window, starting 5’, with a mean quality <Q20, and removed if the final length was less than 35bp. Only paired good quality reads were used downstream. Filtered reads were mapped to SB001 using BWA-MEM v0.7.17 (Li and Durbin 2009). The -M flag was used to flag secondary reads. Mapped reads were further processed using samtools v1.17 (Li et al. 2009), to identify and retain only properly paired reads, fixmates, and sort the reads by coordinates. Picard Tools v2.27.5 MarkDuplicates (Broad Institute, n.d.) was used to flag duplicated reads. Variant calling in each sample was performed using Freebayes v1.3.6 (Garrison and Marth 2012) using the following parameters --min-mapping-quality 30 --min-base-quality 30 --ploidy 1 --min-coverage 50 on individual samples. Comparison of SB001 and CBS8550 was carried out using BCFtools (part of samtools), ‘bcftools stats’ was used to compare and output information and ‘bcftools isec’ was used to generate VCF files containing variants shared by and unique to SB001 vs CBS8550 assuming position and variant are exactly the same in each comparison. A summary of results was created and annotated with the closest two genes to each variant using RnaChipIntegrator (Briggs, Donaldson, and Zeef, n.d.) using the parameters ‘--cutoff=10000 --edge=both --number=2 --compact’. A summary of InDel lengths was created using vcftools v0.1.16 (Danecek et al. 2011) using the parameter ‘--histindel-len’.

Effects of short variants unique to SB001 and CBS8550 were analysed using SnpEff (v4.3T) (Cingolani et al., 2012) for variant annotation and effect prediction. The SnpEff database for *S. batistae* was constructed using the genome sequence and annotation files. To verify, genes containing unique variants in SB001 and CBS8550 were amplified by PCR and sanger sequenced with the Eurofins Mix2Seq service (Eurofins Scientific, Luxembourg). Forward and reverse sequences were aligned in MEGA 11 to build a consensus sequence. Function of genes unique to *S. batistae* were then predicted using Alphafold (Jumper et al., 2021; Varadi et al., 2023) structure prediction and subsequently used as an input for DeepFRI function prediction (Gligorijević et al., 2021).

### Genome assembly

PacBio sequencing data was processed to generate circular consensus sequencing (CCS, or HiFi) reads using the CCS application in SMRT Link 8.0 software package with three passes, considered to generate a minimum Q20 accuracy and to mitigate homopolymer frameshifts (Eid et al., 2009, Sacristan-Horcajada et al., 2021, Wenger et al., 2019). The CCS read length ranged from 333 to 22,100. CCS reads were assembled using the PacBio assembler algorithm’s Improved Phased Assembly (IPA v1.8.0) method (available at https://github.com/PacificBiosciences/pbipa.git). To confirm that our strain is a haploid, pbmm2 v1.10.0 (available at https://github.com/PacificBiosciences/pbmm2), a SMRT C++ wrapper for minimap2’s C API, was used to index the reference genomes and align the sequencing reads to the references. SAMtools v1.10 suite has been used to process the sequence alignment files. BAM files were sorted and indexed for SNP calling using samtools sort and samtools index, respectively. DeepVariant v1.5.073 was used for variant calling. BCFTools v1.10.2 was used for manipulating VCFs and BCFs.

### qPCR validation of mating type

Total RNA was extracted from exponential growth with OD_600_ 0.5 and 0.7 using the RNeasy Mini Kit (Qiagen, Germany) following the manufacturer’s protocol. Residual DNA was degraded with RNase-free DNase set (Qiagen, Germany) according to the manufacturer’s instructions. The extracted RNA was quantified using a NanoDrop Lite Spectrophotometer (Thermo Fisher Scientific, United States). cDNA was reverse transcribed using Go-Script reverse transcriptase (Promega, USA) following the manufacturer’s protocol. qPCR reactions consisted of 2ng of cDNA, 3pmol of each primer, and 5 µl of iTaq Universal SYBR Green Supermix (Bio-Rad, USA) 2X in a final volume of 10 µl. Reactions were carried out on a run on a LightCycler 480 System (Roche, Switzerland) for 35 cycles with the following conditions: 15 s at 95 °C, 30 s at 55 °C, and 30 s at 72 °C. The expression of *STE2*/*STE3* genes were compared against expression of *ACT1*. The primers used in this study are reported in Supplementary Table 10.

### Structural and functional annotation

Telomeres were identified by searching for repeated sequences in the genome assembly using the Telomere Identification toolKit (Brown *et al.,* 2023). Gene annotation and gene prediction was achieved using The Yeast Genome Annotation Pipeline (YGAP) (Proux-Wera et al., 2012) for the *de novo S. batistae* genome assembly. YGAP uses homology and synteny information from other yeast species present in the Yeast Gene Order Browser database to predict the gene structure (based on the hypothesis that the genes intron/exon structure is conserved through evolution). The HybridMine tool v4.0 (Timouma et al., 2020), initially developed for functional annotation at gene level was modified to work at protein level and used to identify one-to-one orthologs between *S. batistae* and *S. cerevisiae*, *Y. lipolytica*, *S. pombe*, *C. albicans, C. glabrata, M. bulderi, M. exigua, K. marxianus, K. lactis*, *M. barnetti, W. sorbophila, S. ingens* and *S. lignohabitans, S. bombicola, S. davenportii, S. stellata, S. bacillaris, S. floricola, S. sirachiensis, S. apicola, and S. vitae*. Gene enrichment analysis was performed on the proteins uniquely shared between *S. batistae* and *S. pombe* using AnGeLi (Bitton et al., 2015). HybridMine was also used to identify groups of homologs within the *S. batistae* SB001 genome. Visualization of the proteins shared between the different species was carried out using the “ComplexUpset” and “ggplot2” R packages. Core genome analysis with “ClusterProfile” R package and “gProfiler”.

### Comparative genome analysis

Evolutionary analysis was conducted in MEGA11 v11.0.1179. This analysis involved 14 species and 2019 conserved protein coding genes. The evolutionary history was inferred by using the Maximum Likelihood method and Tamura-Nei model on the concatenated alignment of the 2019 conserved genes. The tree with the highest log likelihood was selected. Initial tree(s) for the heuristic search were obtained automatically by applying Neighbor-Join and BioNJ algorithms to a matrix of pairwise distances estimated using the Tamura-Nei model and then selecting the topology with superior log likelihood value. Orthologous proteins were first aligned using the MUSCLE global alignment tool, resulting in 2019 alignments. These alignments were then concatenated into a single alignment. The phylogenetic tree was reconstructed using the Minimum Evolution method with a Poisson model. All *Starmerella* genome assemblies were taken from NCBI (Supplementary Table 5). Enrichment analysis of the genes uniquely shared between *Sz. pombe* and *S. batistae* were performed using FYPO (Harris et al., 2013). Figures were made using Python 3.10.

### DAPI staining and mapping of mtDNA

DNA visualisation was performed via DAPI (4,6-diamidino-2-phenylindole; Sigma) staining. Yeast cells were harvested after overnight growth in YP[+[2% glucose and washed twice with PBS. Cells were resuspended in SD media w/o amino acids. DAPI and SDS were added to the culture at the final concentration of 1 µg/ml and 0.01%, respectively. The cells were incubated in the dark for 10 min at 30 °C. Cells were observed with an Eclipse TE2000-U fluorescence inverted microscope (Nikon) fitted with a ×100 immersion objective. The images were captured using the Ocular Image Acquisition Software V2.0 (QImaging). The images were then processed and assembled with Image J.

## Acknowledgments

The authors wish to thank the Genomic Technologies and Bioinformatic Core Facility at the University of Manchester and Tanda Qi for assisting with the SNP variant effect analysis.

## Data availability

The annotation of *Starmerella* yeasts is available at https://github.com/Sookie-S/Starmerella_species_annotation/tree/main.

