## Supplementary figures and images for "Construction of a reference genome for *Starmerella batistae* and annotation of *Starmerella* species reveal a close evolutionary relationship with *Schizosaccharomyces pombe* and suggest an alternative pathway for sophorolipid production"

### Supplementary Figure 1

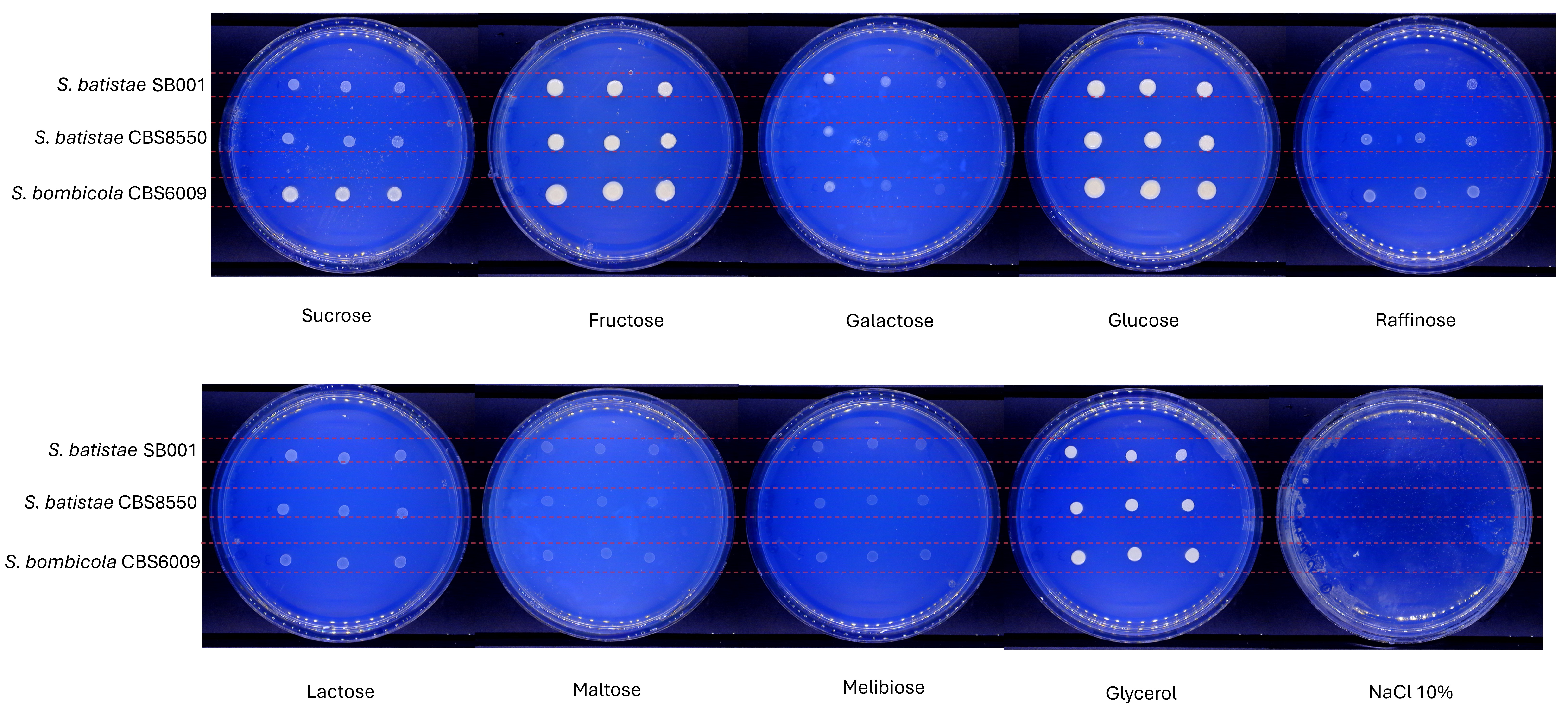

### Supplementary Figure 2

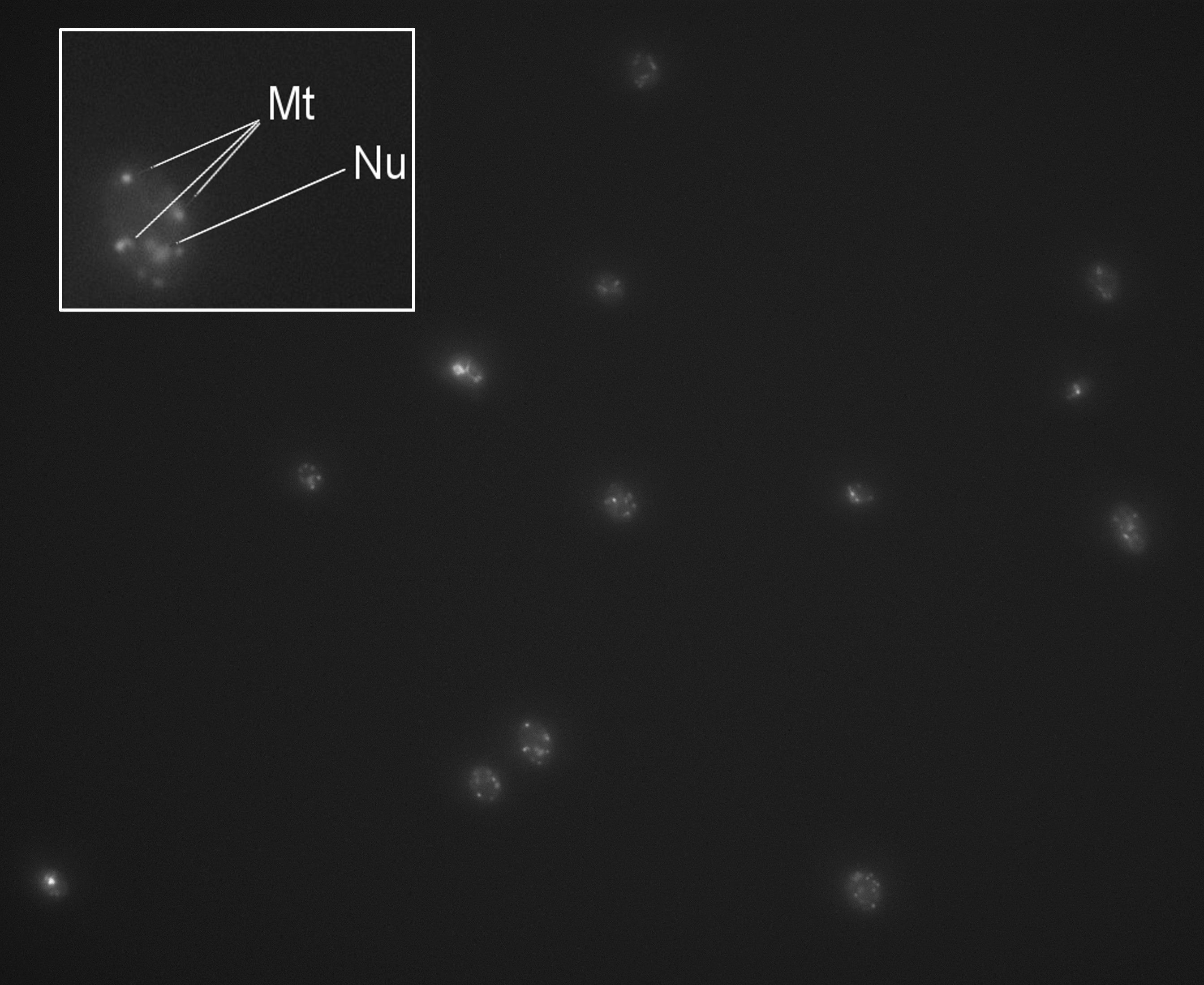

### Supplementary Figure 3

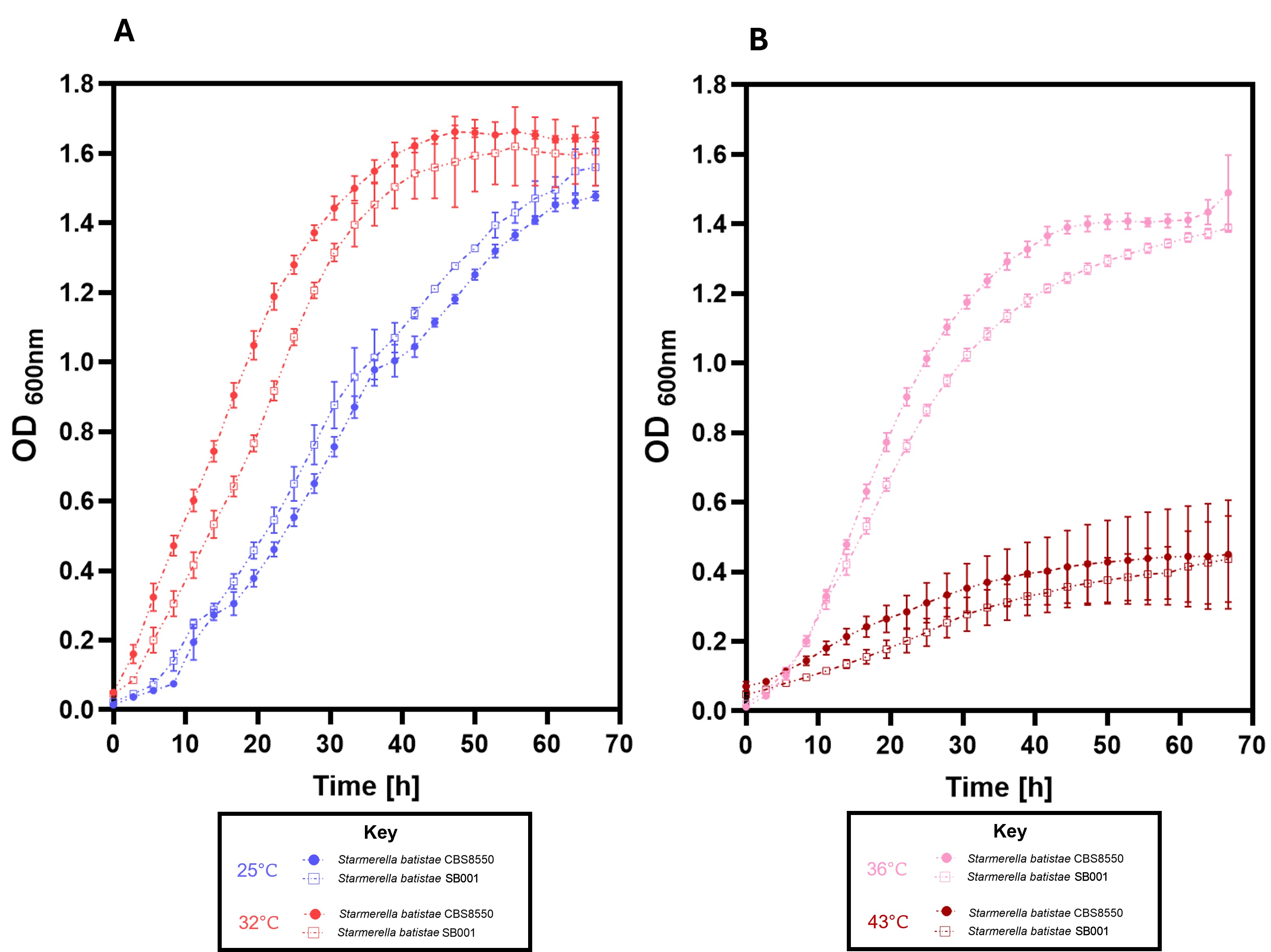

### Supplementary Figure 4

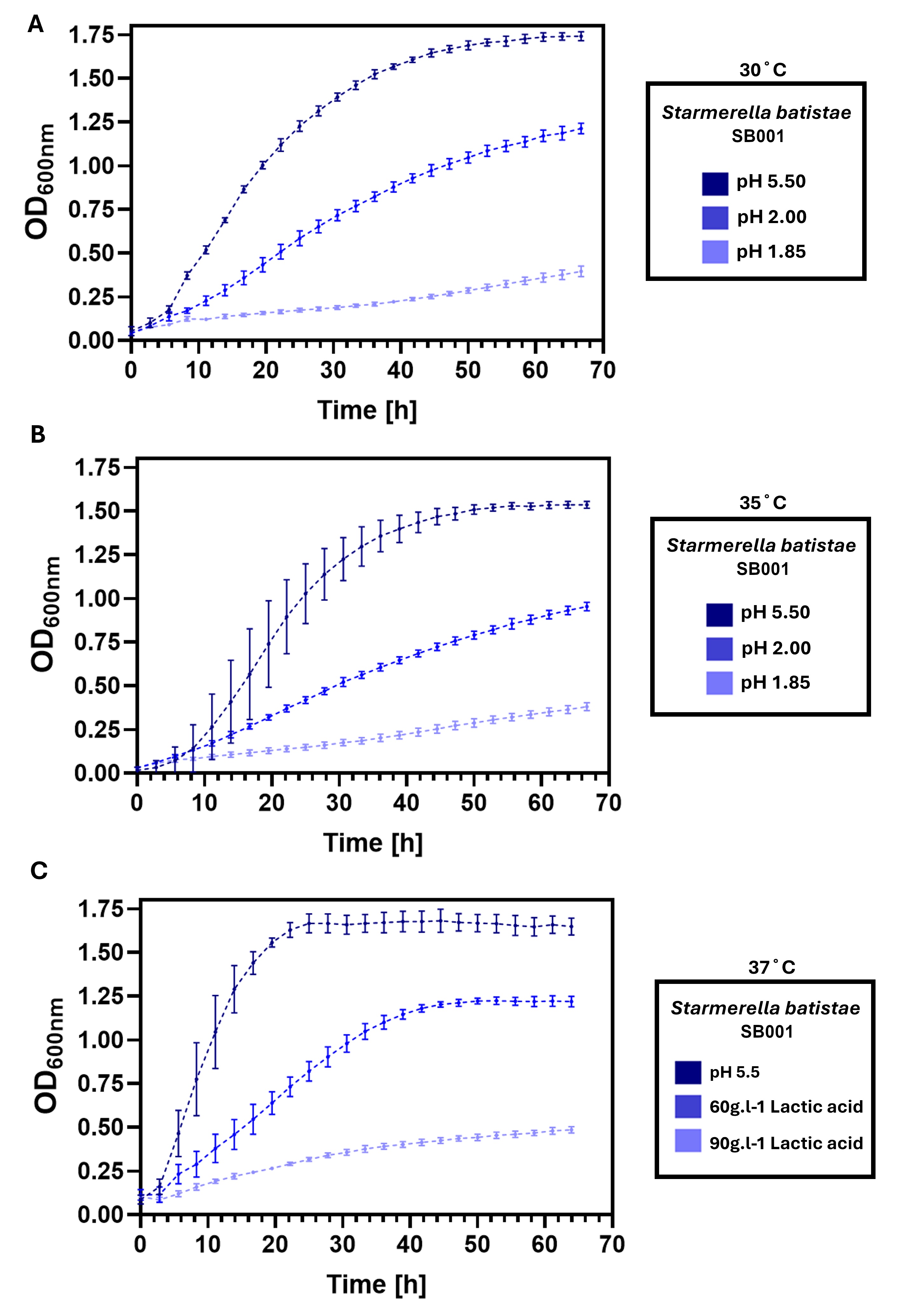

### Supplementary Figure 5

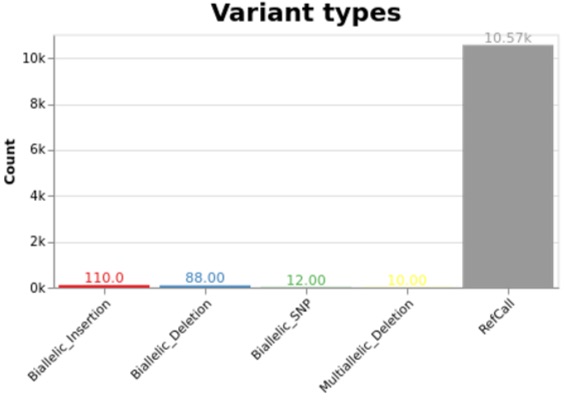

### Supplementary Figure 6

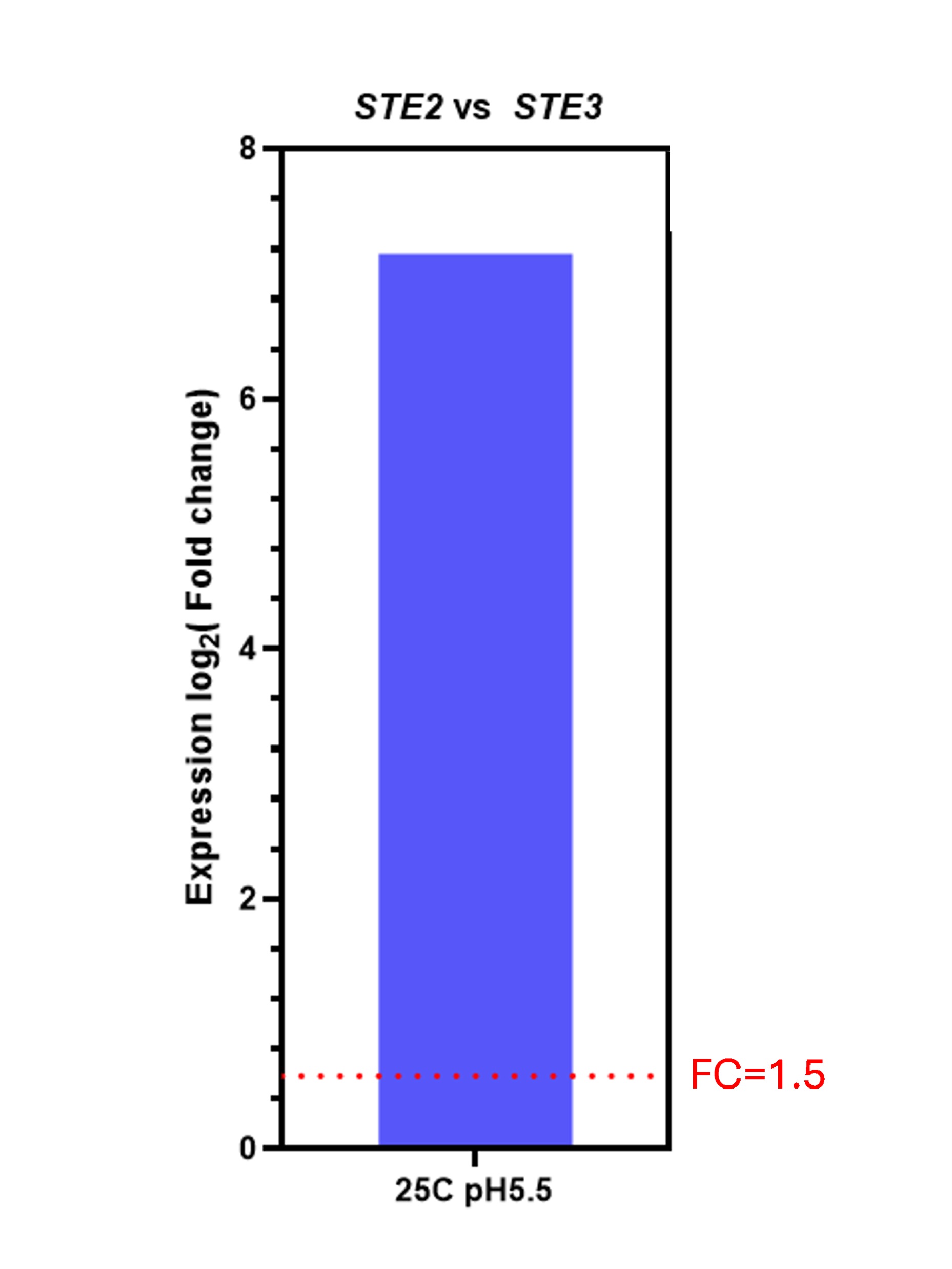

### Supplementary Figure 7

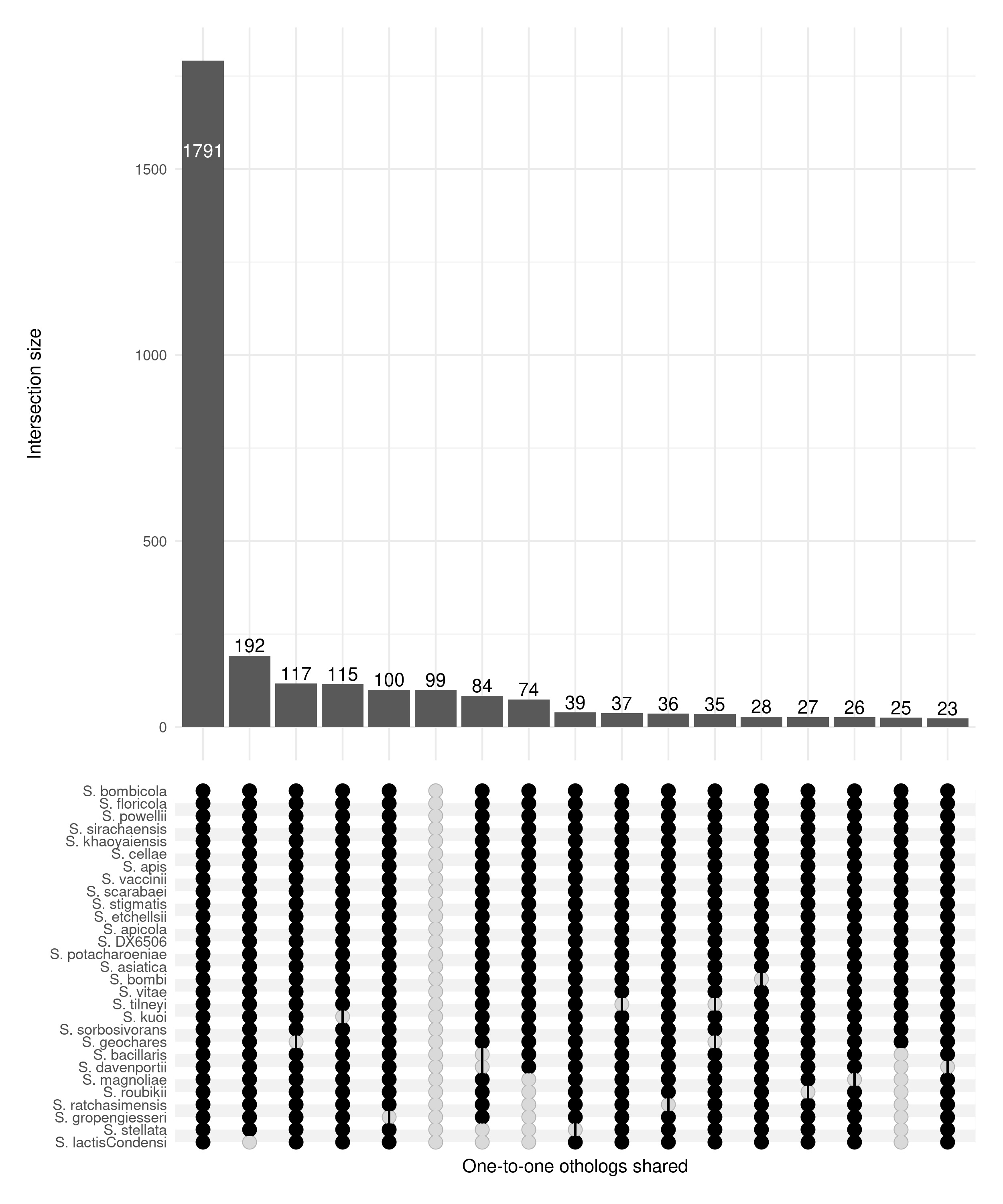

### Supplementary Figure 8

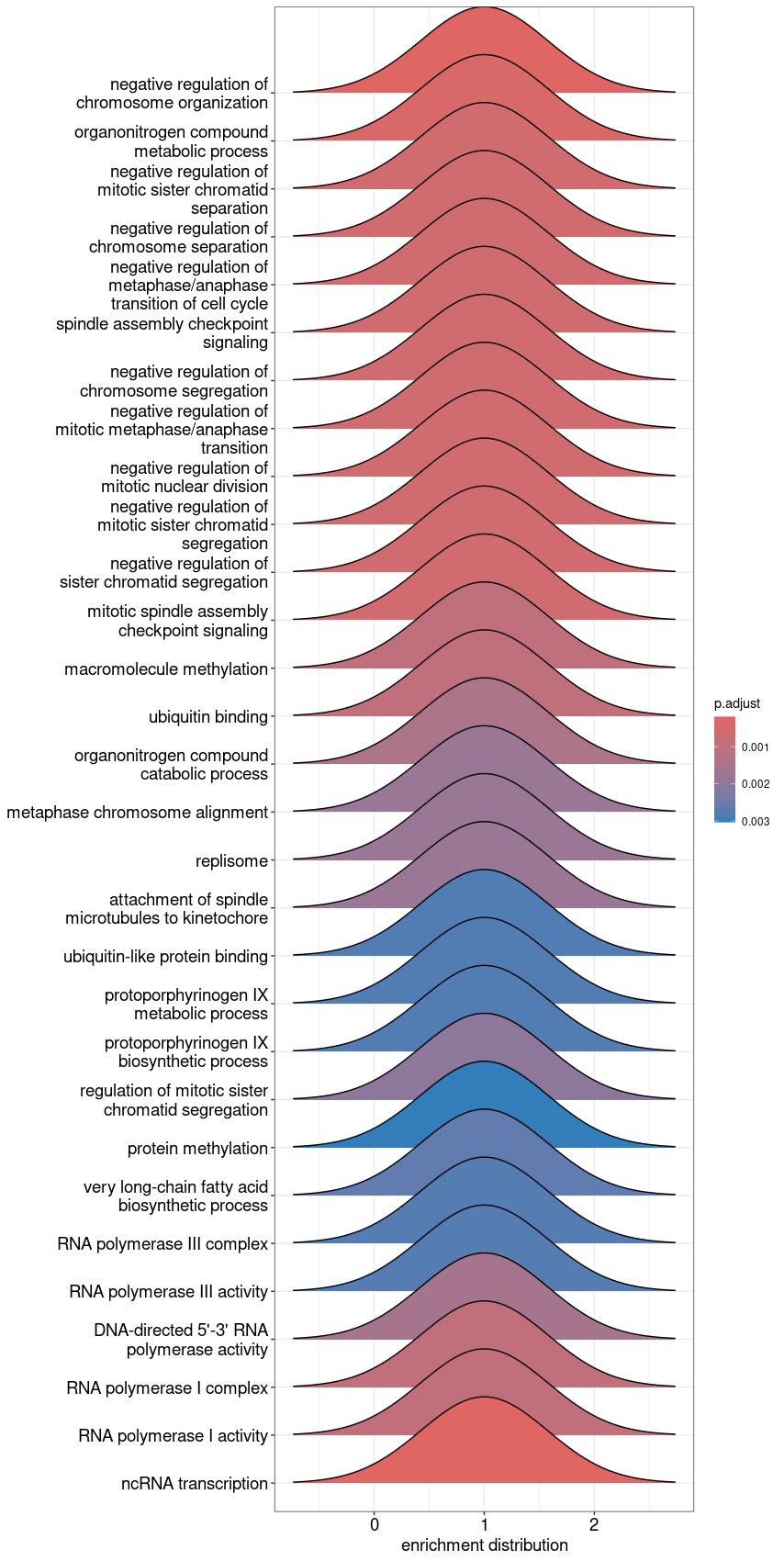

### Supplementary Figure 9

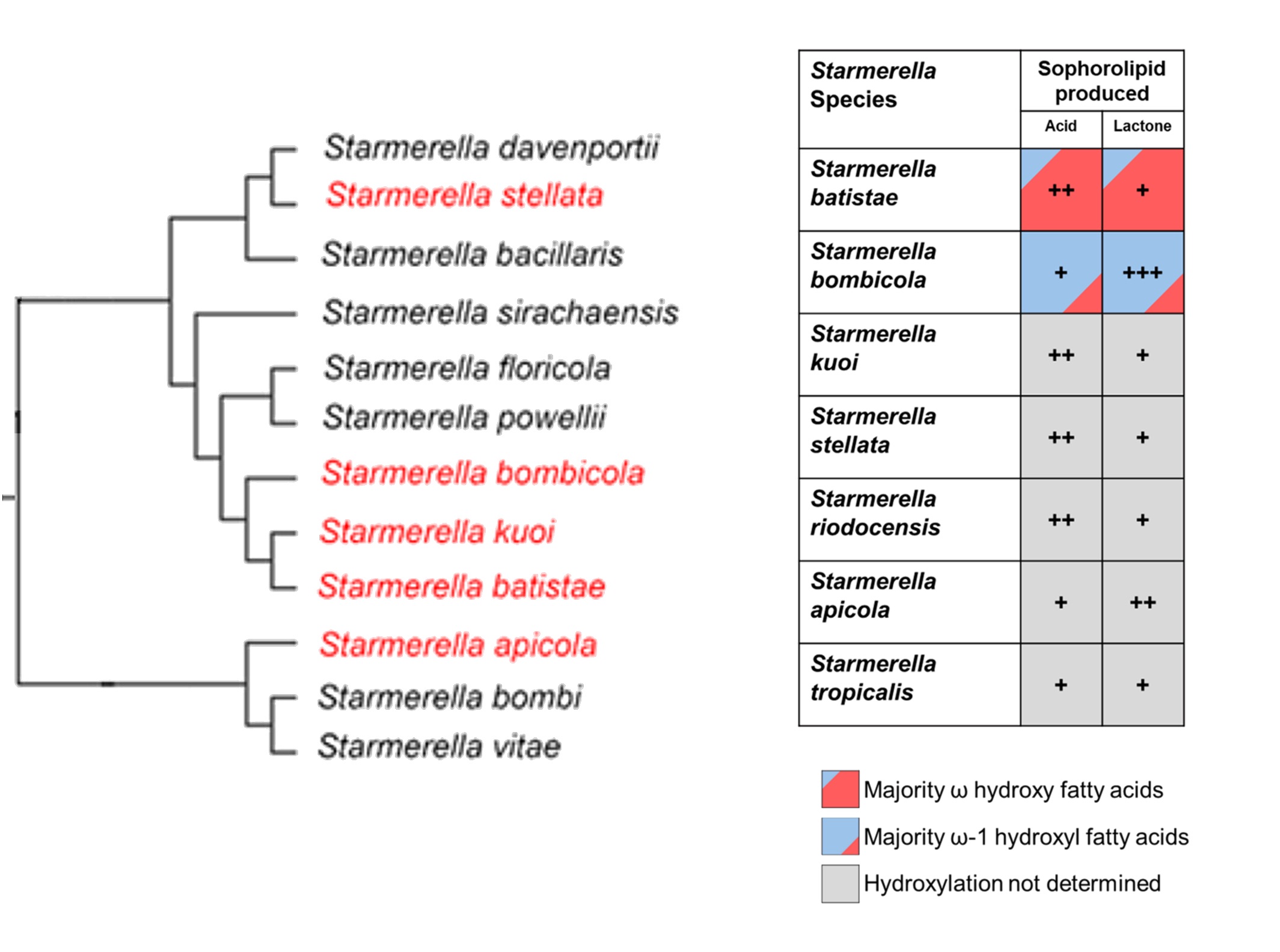
